# CaMK (CMK-1) and O-GlcNAc transferase (OGT-1) modulate mechanosensory responding and habituation in an interstimulus interval-dependent manner in *Caenorhabditis elegans*

**DOI:** 10.1101/115972

**Authors:** Evan L. Ardiel, Troy A. McDiarmid, Tiffany A. Timbers, Kirsten C. Y. Lee, Javad Safaei, Steven L. Pelech, Catharine H. Rankin

**Affiliations:** Djavad Mowafaghian Centre for Brain Health, University of British Columbia, 2211 Wesbrook Mall, Vancouver, British Columbia, V6T 2B5 Canada; Department of Psychology, University of British Columbia, 2136 West Mall, Vancouver, British Columbia, V6T 1Z4 Canada; Department of Computer Science, University of British Columbia, 2366 Main Mall, Vancouver, British Columbia, V6T 1Z4 Canada; Department of Medicine, University of British Columbia, 2775 Laurel Street, Vancouver, British Columbia, V5Z 1M9 Canada, Kinexus Bioinformatics Corporation, Suite 1, 8755 Ash Street, Vancouver, British Columbia, V6P 6T3 Canada

**Author notes:** Co-first authors (Evan L. Ardiel, Troy A. McDiarmid, Tiffany A. Timbers). Correspondence concerning this article should be addressed to: Catharine H. Rankin, Djavad Mowafaghian Center for Brain Health and Department of Psychology, University of British Columbia, 2136 West Mall, Vancouver, British Columbia, Canada, V6T 1Z4.

**Keywords:** Learning, O-GlcNAc, OGT, CaMK1, CaMK4, CMK-1, *C. elegans*, Non-associative learning, Habituation, Mechanosensation, Calmodulin-dependent kinase

## Abstract

The ability to learn is an evolutionarily conserved adaptation that remains incompletely understood. Genetically tractable model organisms facilitate mechanistic explanations of learning that span genetic, neural circuit, and behavioural levels. Many aspects of neural physiology, including processes that underlie learning (*e.g.* neurotransmitter release and long-lasting changes in synaptic strength), are regulated by brief and local changes in [*μ*m] levels of free intracellular Ca^2+^. On this scale, changes in [Ca^2+^] activate many Ca^2+^-sensors, including the Ca^2+^/calmodulin-dependent kinases (CaMKs). Here we reveal that the *Caenorhabditis elegans* ortholog of CaMK1/4, CMK-1, functions in primary sensory neurons to regulate responses to mechanical stimuli and behavioral plasticity, specifically habituation, a conserved form of non-associative learning. The habituation phenotypes of *cmk-1* mutants were dependent on interstimulus interval (ISI), such that CMK-1 slows habituation at short ISIs, but promotes it at long ISIs. We predicted potential CaMK phosphorylation targets from catalytic site analysis of the human and *C. elegans* CaMKs and mutant analysis of these candidates implicated O-linked N-acetylglucosamine (O-GlcNAc) transferase, OGT-1, in mechanosensitivity and learning. Cell specific rescue and knockdown experiments showed that both CMK-1 and OGT-1 function cell autonomously in mechanosensory neurons to modulate learning. Interestingly, despite their similar mutant phenotypes, detailed behavioral analysis of double mutants demonstrated that CMK-1 and OGT-1 act in parallel genetic pathways. Our research identifies CMK-1 and OGT-1 as co-expressed yet independent regulators of mechanosensitivity and learning.

## Introduction

Learning is an evolutionarily conserved process by which organisms update and modify their behavioral output based on past experience. This important phenomenon facilitates survival in constantly changing environments. Habituation is a conserved form of non-associative learning thought to be essential for more complex behavioral plasticity and normal cognitive function (McDiarmid, Bernardos, and Rankin 2017). It is commonly observed as a behavioural response decrement following repeated stimulation that cannot be explained by sensory adaptation or motor fatigue (Catharine H. Rankin et al. 2009). For a comprehensive account of learning from genes to circuits to behvaiours, many researchers have turned to genetically tractable model organisms with well-characterized nervous systems, such as the nematode *Caenorhabditis elegans* (Ardiel and Rankin 2010; McDiarmid, Yu, and Rankin 2017).

*C. elegans* exhibits many forms of behavioural plasticity, including habituation. The *C. elegans* reversal response to repeated non-localized mechanosensory stimuli (taps to the side of the Petri plate) habituates, which can be observed as a decrease in both the size of the reversal (response magnitude) and the likelihood of responding (response probability) (Rankin and Broster 1992; Beck and Rankin 1993). A defining characteristic of habituation is that responses decrement faster to stimuli delivered with a short interstimulus interval (ISI) than to stimuli delivered with a long ISI. Parametric analysis of tap habituation in *C. elegans* led Rankin and Broster (C. H. Rankin and Broster 1992) to propose that multiple distinct mechanisms underlie habituation at different ISIs, but the molecular components of the ISI-dependent mechanism have not yet been delineated.

Ca^2+^ is an important modulator of tap habituation in *C. elegans* (Kindt et al. 2007; Suzuki et al. 2003). Repeated mechanical stimulation results in an attenuation of the associated Ca^2+^ transient (Suzuki et al. 2003) and Ca^2+^ chelation in the mechanosensors results in more rapid habituation, as do mutations in the genes encoding calcium signaling molecules calreticulin (*crt-1*) and the inositol triphosphate receptor (*itr-1:* (Kindt et al. 2007)). Despite the importance of Ca^2+^ signaling in tap habituation, key second messengers by which Ca^2+^ exerts its control over habituation remain unknown. Brief and local changes in micromolar concentration levels of free intracellular Ca^2+^ are known to activate many Ca^2+^-sensors, including ubiquitously expressed calmodulin (CaM). Once bound with four Ca^2+^ ions, CaM (Ca^2+^/CaM) regulates many signaling proteins, including the CaM-kinase (CaMK) family of protein-serine/threonine kinases, which are highly expressed in the nervous system. Within the larger CaMK group (consists of ∼ 23 kinase families) the CaMK1 family (consisting of CaMKK, CaMK1 and CaMK4; reviewed in Soderling 1999) has been shown to play important roles in nervous system development and plasticity. In the developing mammalian nervous system, CaMK1 regulates axonal growth cone motility and axonal outgrowth (Wayman et al. 2004), dendritic arborization (Wayman et al. 2006; Takemoto-Kimura et al. 2007; Wayman et al. 2008), and formation of dendritic spines and synapses (Saneyoshi et al. 2008). In terms of nervous system plasticity, Schmitt et al. (Schmitt et al. 2005) demonstrated that CaMKK and CaMK1, but not CaMK4, activate Ras-extracellular signal-regulated protein kinase (Ras-ERK) signaling during early-phase long-term potentiation (LTP) in hippocampal neuron cultures. Using the same experimental preparation, Guire *et al*. (Guire et al. 2008) found that CaMK1 functions during LTP to recruit calcium permeable AMPA receptors. The *C. elegans* genome encodes a single homolog of CaMK1/4, CMK-1 (Eto et al. 1999). Strains with mutations in *cmk-1* are superficially wild-type (Kimura et al. 2002), however a recent series of studies have implicated CMK-1 in several forms of behavioural plasticity, including experience-dependent thermotaxis and heat avoidance (Yu et al. 2014; Schild et al. 2014; Kobayashi et al. 2016; Lim et al., 2017).

Here we show that *cmk-1* mutants exhibit age-dependent mechanosensory hyperresponsivity phenotypes and learning deficits. The habituation phenotypes of *cmk-1* mutants are sensitive to ISI, such that *cmk-1* inhibits habituation at short ISIs and promotes it at long ISIs. A screen for predicted targets of CMK-1 led us to the *C. elegans* O-linked N-acetylglucosamine (O-GlcNAc) transferase homolog, OGT-1. Cell specific rescue and knockdown experiments revealed that both CMK-1 and OGT-1 function cell autonomously in the mechanosensory neurons to modulate learning. Interestingly, detailed behavioral analysis of double mutants demonstrated that CMK-1 and OGT-1 act in parallel pathways to influence behavioral plasticity.

## Results

### CMK-1 differentially modulates habituation at long versus short ISIs

Wild-type worms habituate to repeated tap stimuli by decreasing the distance they reverse, the shorter the interval between stimuli, the faster the decrement (Rankin and Broster 1992; Beck and Rankin 1993). To determine whether CaMK1/CaMK4 functions in learning, we examined habituation of the *C. elegans* putative null mutant *cmk-1(oy21*). At a 10s ISI, the *cmk-1* mutants habituated more rapidly than wild-type to a similar asymptotic level (Fig. 1A). That is, neither the initial nor final response size of *cmk-1* mutants differed from wild-type, but the rate of decline increased, as determined by the time constant (τ) of a two-term exponential fit of the habituation curves (Fig. 1B). In contrast, when the rate of tapping was slowed to a 60s interval (Fig. 1C), *cmk-1* mutants displayed the opposite deficit, i.e. reduced habituation to a higher asymptotic level (Fig. 1D). Importantly, the *cmk-1* habituation deficit at both ISIs could be rescued by expression of CMK-1 cDNA from the endogenous promoter (Fig. 1), thereby confirming *oy21* as the causative allele.

**Figure 1.**
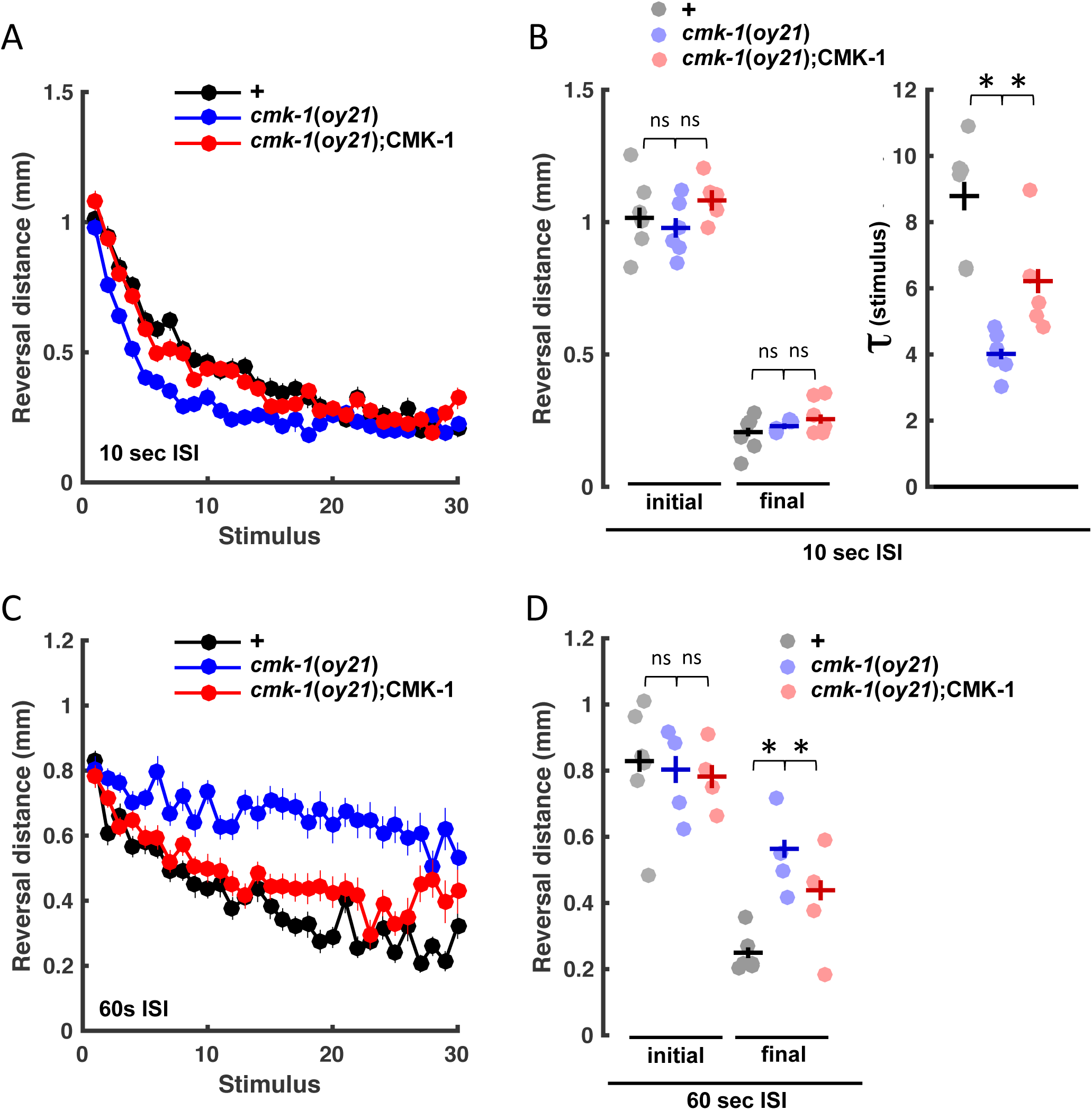
CMK-1 modulates tap habituation in an ISI-dependent manner. (A) Length of reversal response to 30 taps administered at a 10s ISI. Mean +/- SEM. (B) Initial response length and final response level for individual plates with population mean +/- SEM from ‘A’ (left). Individual plate time constants (*τ*) and mean +/- SEM for data in ‘A’ (right). (C) Length of reversal response to 30 taps administered at a 60s ISI. Mean +/- SEM. (D) Initial response length and final response level for individual plates with population mean +/- SEM from ‘C’. Asterisks denote *p*<0.05 versus ‘ns’ for *p*>0.05.

### Age-dependent mechanoresponsivity

As worms develop from 72-120 h old (i.e. day 1 adults to day 3 adults), they exhibit deficits in mechanosensory intensity discrimination that are readily apparent in habituation assays, but not necessarily naïve responses (Timbers et al. 2013). We found that *cmk-1* mutants developed a naive touch response phenotype, such that their reversals were larger than wild-type as day 2 and 3 adults (96h and 120h of age), but not as day 1 adults (72h of age; Fig. 2A). Years of screening have identified several touch insensitive mutants, but loss of *cmk-1* causes a rare hyper-responsive phenotype, i.e. larger reversal responses than wild-type. To investigate how hyper-responsivity affected responding to repeated stimulation, day 2 adults were given taps at a 10s or 60s ISI. If stimuli were presented every 10 s, the *cmk-1* mutant habituated faster than wild-type to an asymptotic level indistinguishable from wild-type (Fig. 2B & 2C). When taps were administered at a 60s ISI, the large response of cmk-1 relative to wild-type persisted across the training session (Fig. 2D & 2E). Thus, the final level of responding relative to wild-type was dependent on the rate at which stimuli were delivered, suggesting that initial and final response size are dissociable. This assertion was unexpectedly verified with a *cmk-1* presumed gain-of-function allele, *pg58*, which has also been shown to share features with *cmk-1* loss-of-function alleles. Isolated by Schild et al. (2014) in a screen for noxious heat avoidance defects, *cmk-1*(*pg58*) encodes a truncated protein lacking most of its regulatory domain and a nuclear export sequence (NES), but with an intact kinase catalytic domain. As had been observed with a similar mutation in mammalian CaMKI (Stedman et al. 2004), Schild et al. (2014) found that truncated CMK-1(1-304) abnormally accumulated in the nucleus. As with day 2 *cmk-1*(*oy21*) mutants, the *pg58* allele resulted in a large initial response to tap, but the final habituated response size of *cmk-1*(*pg58*) was indistinguishable from wild-type (Fig. 2F). The *pg58* allele therefore genetically dissociates initial and final response phenotypes and suggests that appropriate subcellular localization of CMK-1 is essential for setting naïve responsivity to tap, but not necessarily for modulating it. To simultaneously study habituation and mechanoresponsivity, we focused on 60s ISI tap habituation of day 2 adults, comparing initial response size as a metric for mechanoresponsivity and the final response size as a metric for habituation.

**Figure 2.**
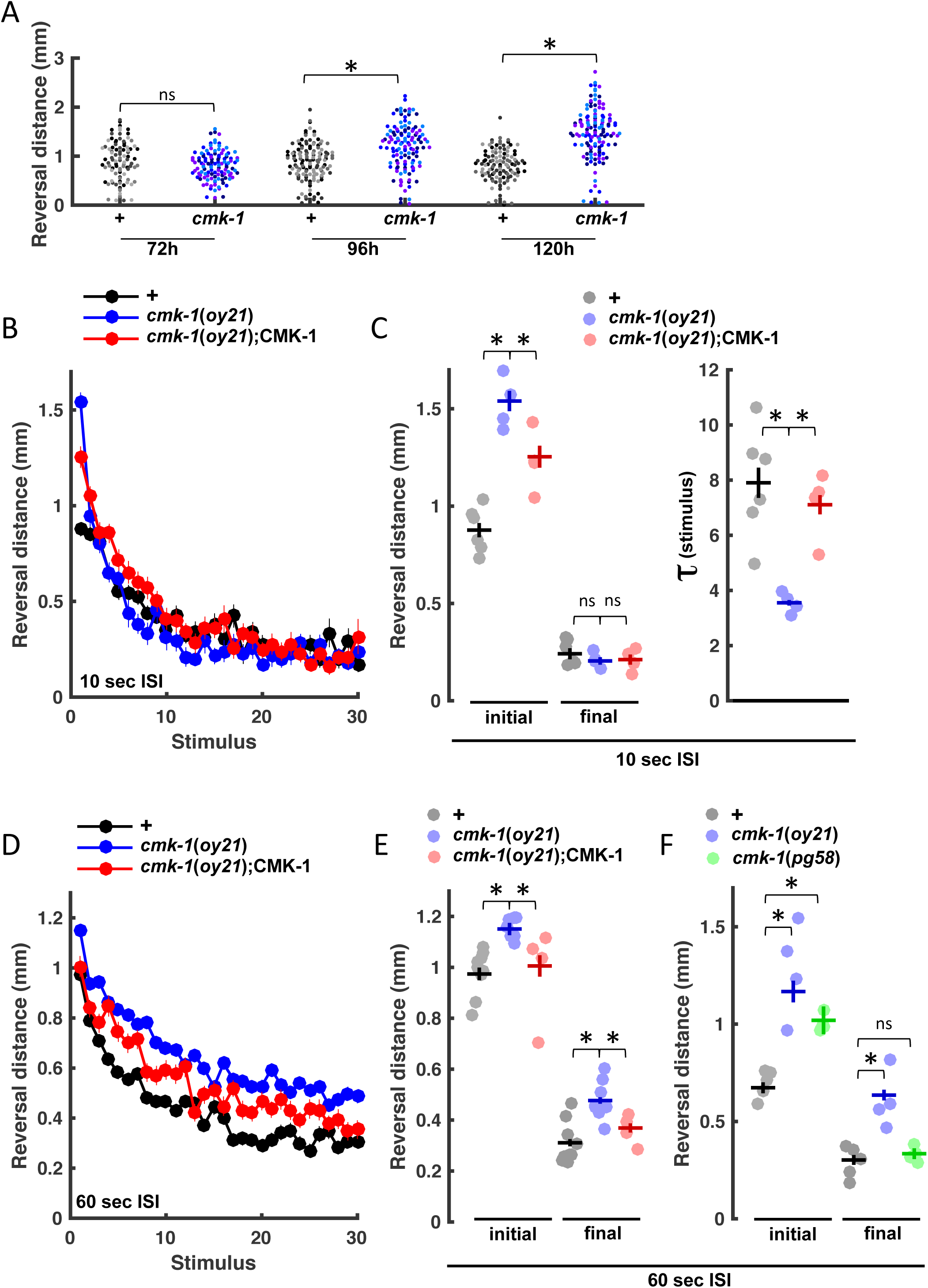
CMK-1 modulates the tap response in an age-dependent manner. (A) Naïve tap response size across adulthood. Markers correspond to individual worms and are color-coded by plate. (B) 96 h old worms tapped at a 10s ISI. Mean +/- SEM. (C) Initial response length and final response level for individual plates with population mean +/- SEM from ‘B’ (left). Individual plate time constants (τ) and mean +/- SEM for data in ‘B’ (right). (D) 96 h old worms tapped at a 60s ISI. Mean +/- SEM. (E) Initial response length and final response level for individual plates with population mean +/- SEM from ‘D’. (F) Initial response length and final response level for individual plates with population mean +/- SEM. Asterisks denote *p*<0.05 versus ‘ns’ for *p*>0.05.

### CMK-1 functions in mechanosensory neurons independent of CKK-1

Analysis of transgenic lines expressing fluorescent transcriptional reporters (P*cmk-1*::GFP) revealed that *cmk-1* is expressed in the touch receptor neurons and interneurons of the tap withdrawal circuit (Fig. 3A & 3B; Wicks and Rankin 1995). To test if CMK-1 functioned in mechanosensory neurons to modulate the tap response, we assessed *cmk-1(oy21)* mutants expressing a wild-type copy of the CMK-1 cDNA from a *mec-3* promoter. The *mec-3* promoter expresses in the touch cells (ALM L/R, AVM, PLM L/R and PVM), as well as mechanosensitive FLP and PVD. Restoring CMK-1 expression in these mechanosensors rescued the hypersentivity and impaired habituation phenotypes of *cmk-1*(*oy21*) null mutants (Fig. 3C). Thus, CMK-1 can function in primary sensory cells to modulate mechanoresponsivity and habituation.

**Figure 3.**
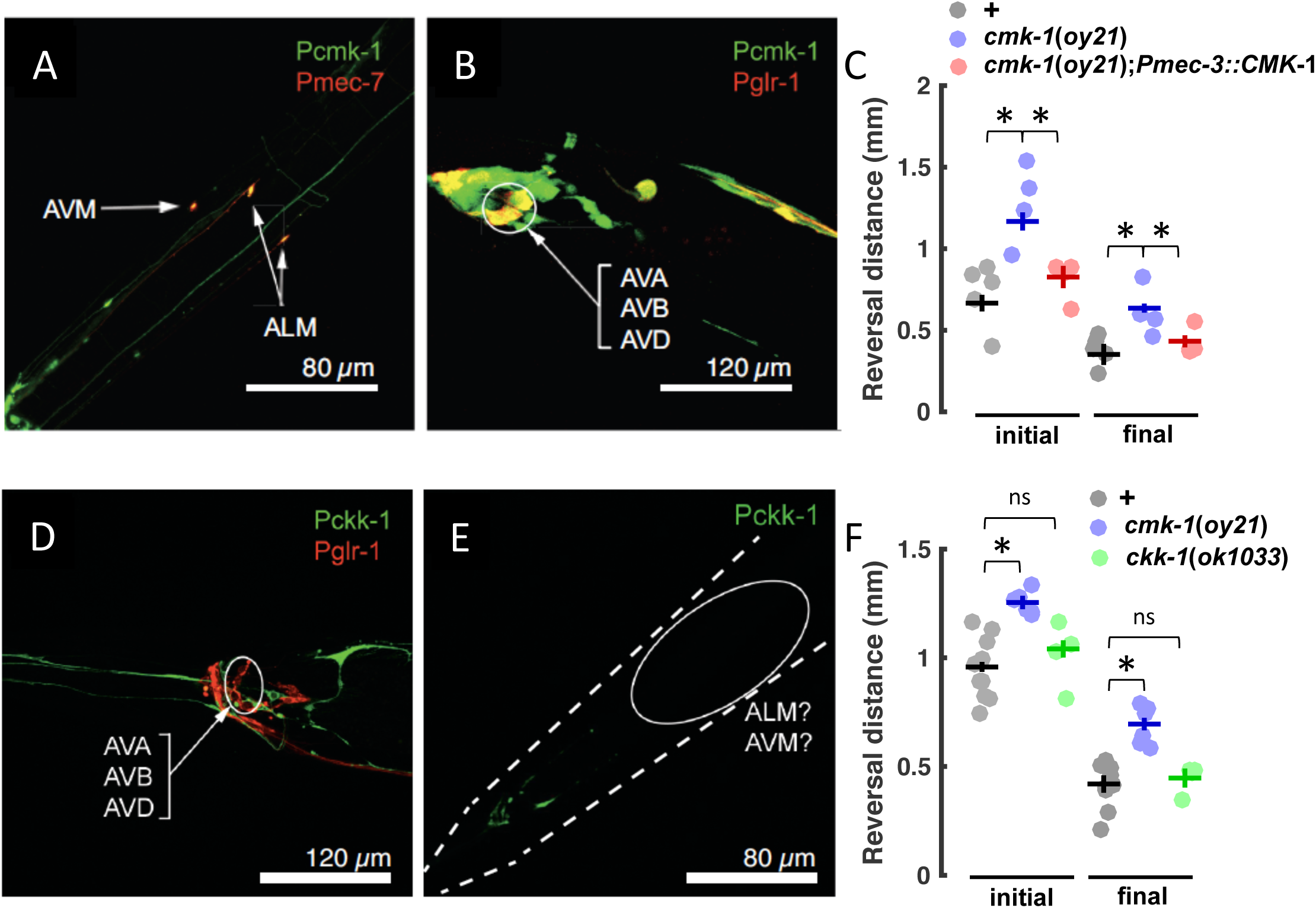
CMK-1, but not CKK-1, functions in mechanosensory neurons. (A) The touch cells (visualized with P*mec-7*::mRFP) express CMK-1 (visualized with P*cmk-1*::GFP). (B) Tap withdrawal interneurons, AVA, AVB, AVD (visualized with P*glr-1*::dsRed) also express CMK-1 (visualized with P*cmk-1*::GFP). (C) Initial response length and final response level for individual plates of 96h old worms tapped at a 60s ISI with population mean +/- SEM. (D) GLR-1 expressing neurons (including the tap withdrawal interneurons, AVA, AVB, AVD; visualized with P*glr-1*::dsRed) do not express CKK-1 (visualized with P*ckk-1*::GFP). (E) No fluorescence (visualized with P*ckk-1*::GFP) was observed where ALM and AVM cell bodies are located (note: a pair of neuronal cell bodies was observed to fluoresce in the tail, but PLM was excluded as a candidate because their neurites projected directly into the ventral nerve cord, whereas PLM projects anteriorly along the lateral sides of the body). (F) Initial response length and final response level for individual plates of 96h old worms tapped at a 60s ISI with population mean +/- SEM. Asterisks denote *p*<0.05 versus ‘ns’ for *p*>0.05.

CaMKK phosphorylates Thr residues in both CaMK1 and CaMK4 to increase Ca^2+^/CaM-dependent phosphotransferase activity (Haribabu et al. 1995; Kitani, Okuno, and Fujisawa 1997). However, *cmk-1* is more broadly expressed than *ckk-1*, the *C. elegans* homolog of CaMKK (Kimura et al. 2002). Indeed, *ckk-1* was not expressed in the sensory or inter-neurons of the tap response circuit (Fig. 3D and 3E) and *ckk-1(ok1033)* mutants did not show the same phenotype as *cmk-1*(*oy21*) in the habituation assay, as their initial and final responses were indistinguishable from wild-type (Fig. 3F). These data suggest that either *(i)* activation of CaMK by calmodulin is sufficient to activate the kinase in the context of some biological signaling, and/or *(ii)* CaMK is activated via some other protein. To test these hypotheses we took advantage of a *cmk-1* point mutant *(gk691866)* from the Million Mutation Project (MMP) collection (Thompson et al. 2013). In this allele, the conserved CaMKK phosphorylation site, Threonine-179 (T179;(Haribabu et al. 1995; Kitani, Okuno, and Fujisawa 1997)), was mutated to isoleucine (I) and therefore could no longer be phosphorylated. After outcrossing the MMP strain, we found that, like *cmk-1* nulls, 96 h old *cmk-1*(*gk691866*) mutants had a larger initial response to tap than wild-type and remained more responsive to repeated stimulation at a 60s ISI (Fig. 4A & 4B). To rule out the potential contribution from a recessive background mutation in the heavily mutagenized MMP strain, we also confirmed that *oy21* and *gk691866* alleles failed to complement (Fig. 4C & 4D). Thus, T179 is an important site for modulation of mechanoresponsivity by CMK-1.

**Figure 4.**
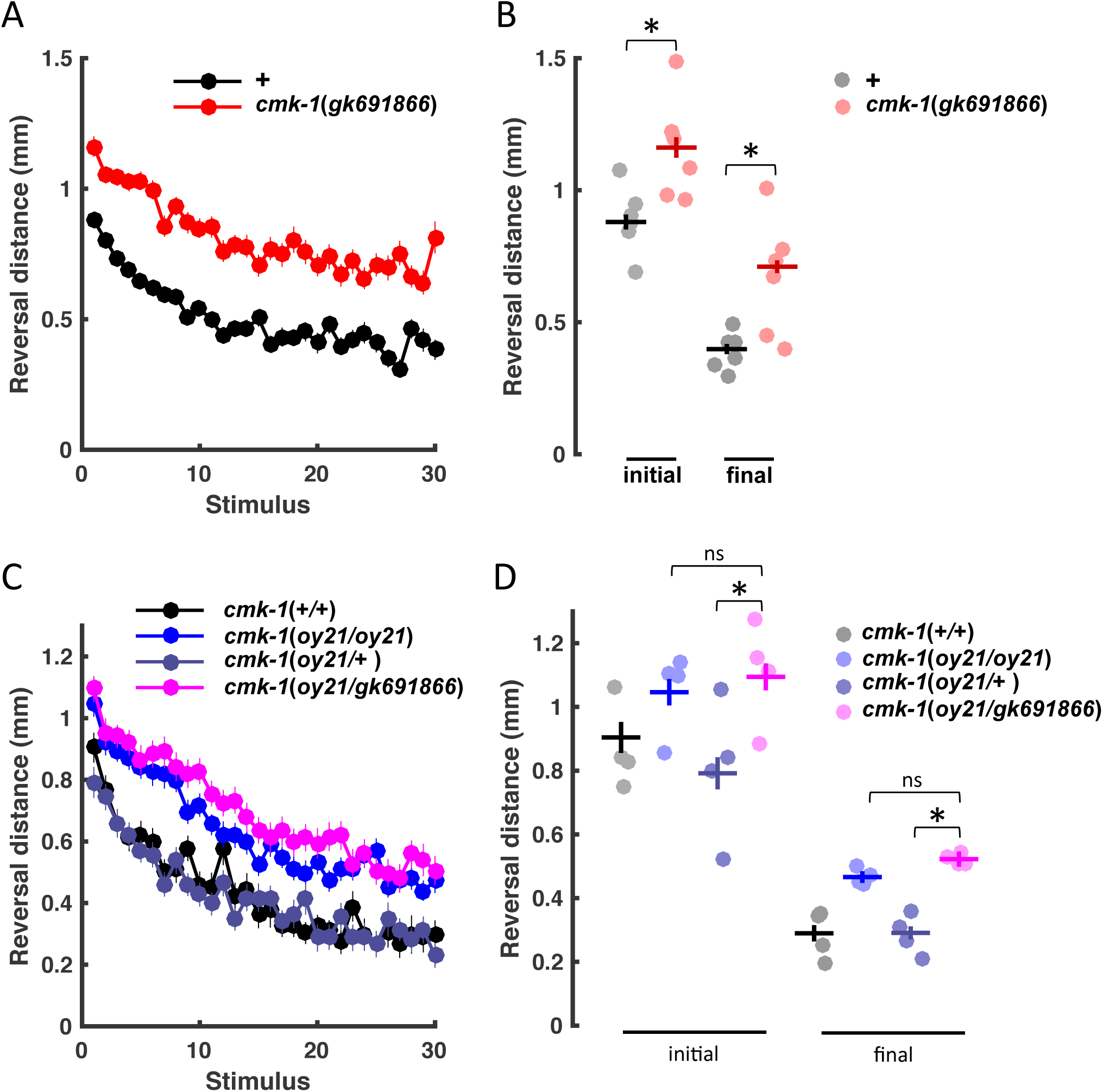
Role of the CMK-1 phosphorylation site (T179). (A) 96h old worms tapped at a 60s ISI. Mean +/- SEM. (B) Initial response length and final response level for Individual plates with population mean +/- SEM used in ‘A’. (C) Complementation test with 96h old worms tapped at a 60s ISI. Mean +/- SEM. (D) Initial response length and final response level for Individual plates with population mean +/- SEM used in ‘C’. Asterisks denote *p*<0.05 versus ‘ns’ for *p*>0.05.

### Identification of evolutionarily conserved predicted CaMK phosphosites

Protein kinases recognize their specific Ser/Thr/Tyr phosphorylation targets based on the sequence of residues that flank the phosphoacceptor site (comprising the kinase consensus sequence). This is akin to a lock and key model, whereby the peptide sequence flanking the phosphosite on the target protein fits into the catalytic domain of the kinase because of the presence of specificity-determining residues that directly interact with the side chains of amino acid sequences surrounding phosphosites (Saunders et al. 2008). These principles have been previously used to predict the kinase substrate specificities of 492 human protein kinases in silico (Safaei et al. 2011). The computational methods have now been further improved with refinements to the original algorithms and training data from over 10,000 kinase-protein phosphosite pairs and 8,000 kinase-peptide phosphosite pairs. We used these updated methods to generate a kinase substrate specificity prediction matrix (KSSPM) for *C. elegans* CMK-1 based on the primary amino acid sequence of its catalytic domain. This KSSPM was used to query all of the 20,470 known *C. elegans* protein sequences. Table S1 lists the top 600 scoring predicted phosphosites. We identified the closest human cognate proteins that featured similar phosphosites, and then scored the human phosphosites with KSSPMs for all four human CaMK1 isoforms and CaMK4 (which share 65% and 44% sequence identity with the *C. elegans* CMK-1 protein, respectively; Fig. S1). Of particular interest were those *C. elegans* proteins and phosphosites that were highly conserved in *Homo sapiens* and predicted to be targeted by human CaMK1 isoforms and CaMK4. Such high evolutionary conservation would support important functional roles for these kinase-substrate pairs. More information about the predicted phosphorylation of these human phosphosites by human protein kinases and their evolutionary conservation in over 20 other species is available in the PhosphoNET website at www.phosphonet.ca.

The list of 600 phosphosites in 373 *C. elegans* proteins was used to prioritize candidates. Proteins that had been previously shown to interact with CaMKs for which testable knockout mutant alleles were available were of highest interest, as well as the most highly ranked by p-site scores. We assayed putative loss-of-function mutant habituation phenotypes for the top candidates. Out of the 22 mutants characterized to date, 17 were observed to show an initial response and/or habituation phenotype (Fig. 5).

**Figure 5.**
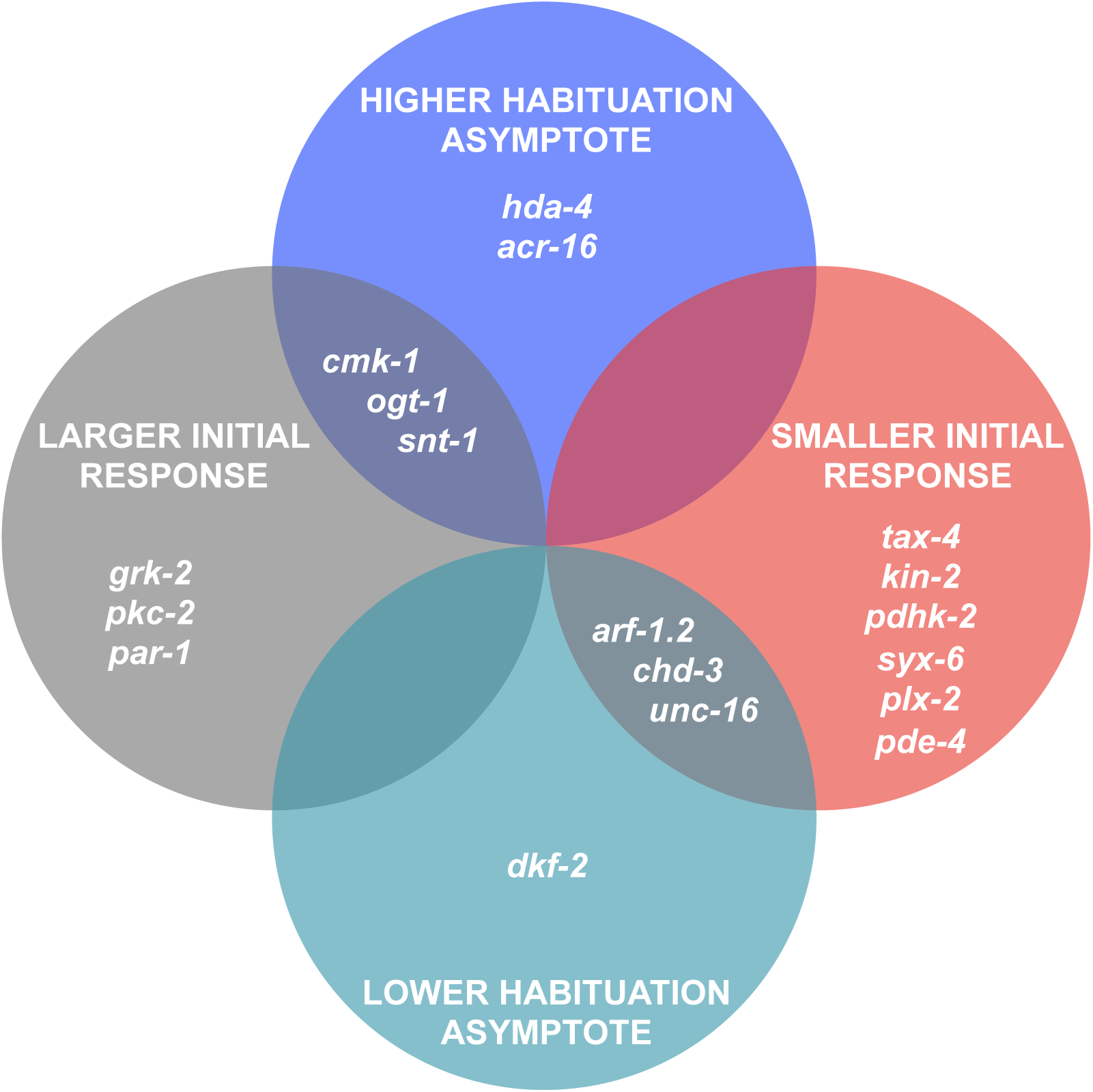
Phenotypes of predicted CMK-1 downstream phosphorylation targets.

### O-GlcNAc transferase, OGT-1, functions in the touch cells to modulate habituation

*ogt-1* mutants showed a mechanosensitivity and habituation phenotype that was strikingly similar to *cmk-1*(*oy21*) mutants, i.e. *ogt-1(ok430)* mutants were significantly more responsive to the initial tap and showed a significantly larger habituated response (Fig. 6A, 6B). We tested whether OGT-1 was also similar to CMK-1 in its ISI dependency. When we habituated the *ogt-1(ok430)* mutant at a 10s ISI instead of a 60s ISI, we observed that they were again more responsive to the initial tap, but habituated more rapidly and to the same final level as wild-type worms in the measure of reversal distance (Fig. 6C). Thus, the *ogt-1* mutants showed the same phenotype as *cmk-1* mutants in all measures tested. Importantly, Hanover *et al*. (Hanover et al. 2005) have demonstrated that the *ogt-1(ok430)* allele results in the complete loss of function of transferase activity. Testing a second null allele of *ogt-1*, *tm1046* (a 466 bp deletion, resulting in a frameshift and an early stop after 392 amino acids), we observed both an initial response and habituation phenotype similar to *ogt-1(ok430)* null mutants (Fig. 6D), thus implicating OGT-1 as a modulator of the tap response.

**Figure 6.**
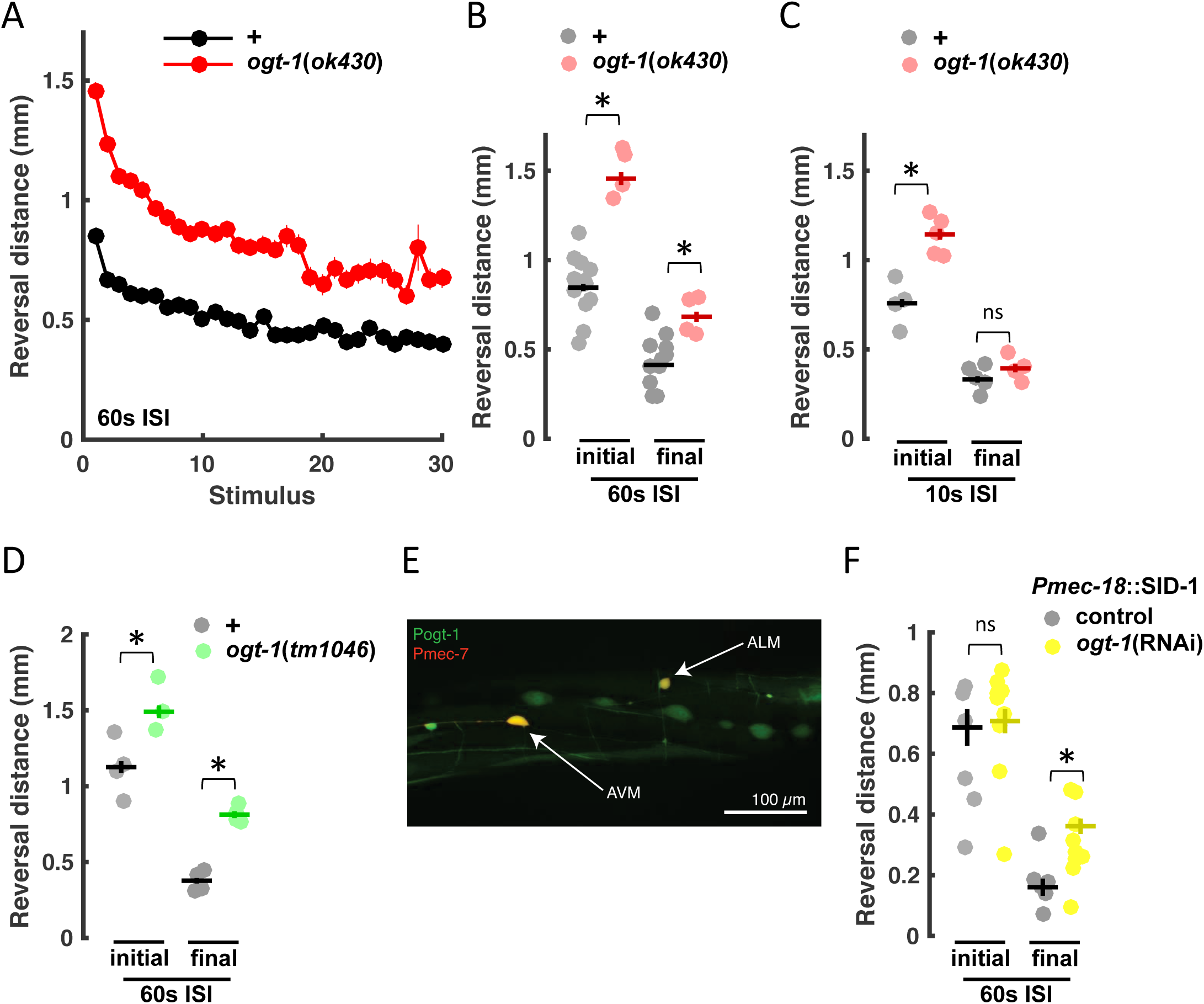
OGT-1 modulates tap responses. (A) 96 h old worms tapped at a 60s ISI. Mean +/- SEM. (B) Initial response length and final response level for individual plates with population mean +/- SEM from ‘A’. (C) Initial response length and final response level for individual plates of 96h old worms tapped at a 10s ISI with population mean +/- SEM. (D) Initial response length and final response level for individual plates of 96h old worms tapped at a 60s ISI with population mean +/- SEM. (E) Mechanosensory neurons, ALMs and AVM (visualized with P*mec-7::mRFP),* express OGT-1 (visualized with P*ogt-1::GFP).* (F) Initial response length and final response level for individual plates of 96h old worms tapped at a 60s ISI with population mean +/- SEM. RNAi by feeding was done in a genetic background targeting dsRNA to the touch cells: TU3568 *sid-1*(*pk3321*) him-5(*e1490*); *lin-15B*(*n744*); *uIs71*[P*myo-2*::mCherry; P*mec-18*::SID-1]. Asterisks denote *p*<0.05 versus ‘ns’ for *p*>0.05.

To evaluate the expression pattern of *ogt-1* we created a transcriptional reporter that consisted of ∼2 kb of the *ogt-1* promoter fused to GFP (P*ogt-1*::GFP). OGT-1 was expressed broadly across the nervous system, including the touch cells (Fig. 6E), in addition to muscles and seam cells. We used cell-specific RNAi by feeding to test whether OGT-1, like CMK-1, functioned in the touch cells. The *C. elegans* nervous system is generally refractory to systemic RNAi, however targeting expression of the dsRNA transporter, SID-1, to neurons creates a dsRNA sink in those cells, allowing for cell specific knockdown when done in a *sid-1* mutant background (Calixto et al. 2010). Using a strain expressing SID-1 exclusively in the touch cells, we showed that RNAi knockdown of *ogt-1* resulted in reduced habituation (Fig. 6F). Taken together, our data demonstrate that OGT-1, like CMK-1, functions cell autonomously in the mechanosensory neurons to influence tap responses.

### *cmk-1* and *ogt*-1 function in parallel pathways

To test whether *cmk-1* and *ogt-1* interact genetically, we assayed the habituation of a *cmk-1(oy21)*; *ogt-1(ok430)* double mutant. The mutant alleles acted additively to give significantly larger naïve and habituated responses than either single mutant (Fig. 7A), suggesting that they function in independent genetic pathways. This result was surprising given the similarity of the individual mutant phenotypes, the common site of action, and the interaction observed in mammalian homologues (Song et al. 2008; Dias et al. 2009). To further probe the double mutant phenotype, we evaluated the behavioral components that determine reversal distance, ie reversal duration and reversal speed. For the initial response to tap, the *cmk-1* single mutant exhibited significantly faster reversals (Fig. 7B) of increased duration (Fig. 7C) compared to wild-type. In contrast, the *ogt-1* mutant reversed at a speed similar to wild-type (Fig. 7B), but with a duration longer than even the *cmk-1* mutant (Fig. 7C). Thus, the single mutant data indicate that the similar reversal distance phenotype of unhabituated *cmk-1* and *ogt-1* mutants was caused by distinct processes, with CMK-1 primarily influencing speed of reversals and modestly influencing reversal duration, and OGT-1 influencing duration, but not speed. However, *cmk-1; ogt-1* double mutants performed significantly slower reversals than *cmk-1* single mutants, almost indistinguishable from wild-type, indicating that *ogt-1* may act as a suppressor of *cmk-1* for reversal speed in naïve animals. Thus, in the context of responding to an initial mechanical stimulus, there may be some interaction between the *cmk-1* and *ogt-1* pathways.In the case of habituated animals, both *cmk-1* and *ogt-1* single mutants exhibited faster reversals of longer duration compared to wild-type. These phenotypes were additive, giving the rapid and long-lasting final responses of the *cmk-1; ogt-1* double mutant.

**Figure 7.**
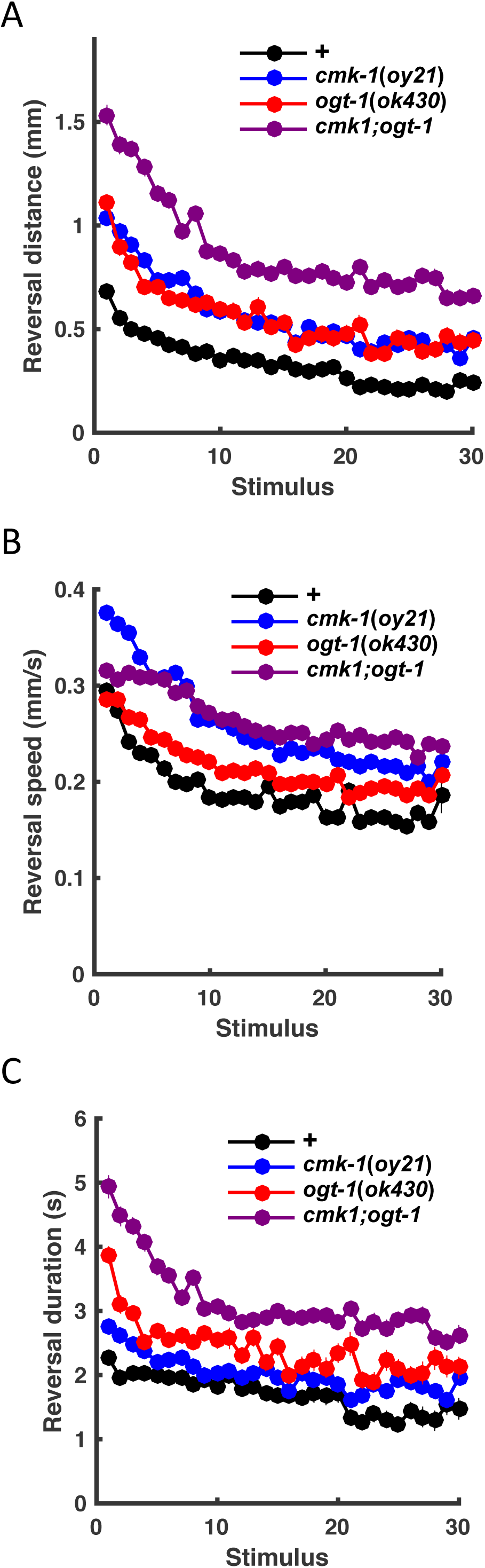
OGT and CMK-1 function in parallel pathways. Reversal distance (A), speed (distance/duration; B), and duration (C) of 96h old worms tapped at a 60s ISI. Mean +/- SEM.

## Discussion

We report here that *C. elegans* CMK-1 functions cell autonomously in mechanosensory neurons to modulate mechanosensitivity and learning in an age- and ISI-dependent manner. We show that *cmk-1* influences this form of behavioural plasticity independent of its canonical upstream activating kinase CKK-1 (Fig. 3F), and that its conserved phosphorylation site at T179 is still required (Fig. 4). Following catalytic site analysis of CaMKs we screened potential phosphorylation targets and implicated OGT-1 in mechanosensory responding and learning. While both CMK-1 and OGT-1 function in mechanosensory neurons to modulate learning (Fig. 3C & 6F), detailed behavioral analysis of *cmk-1(oy21);ogt-1(430)* double mutants demonstrated that they act in parallel genetic pathways to influence behavioral plasticity (Fig. 7 & 8). Our work identifies CMK-1 and OGT-1 as co-expressed yet independent cell autonomous ISI-dependent regulators of mechanosensitivity and learning.

**Figure 8.**
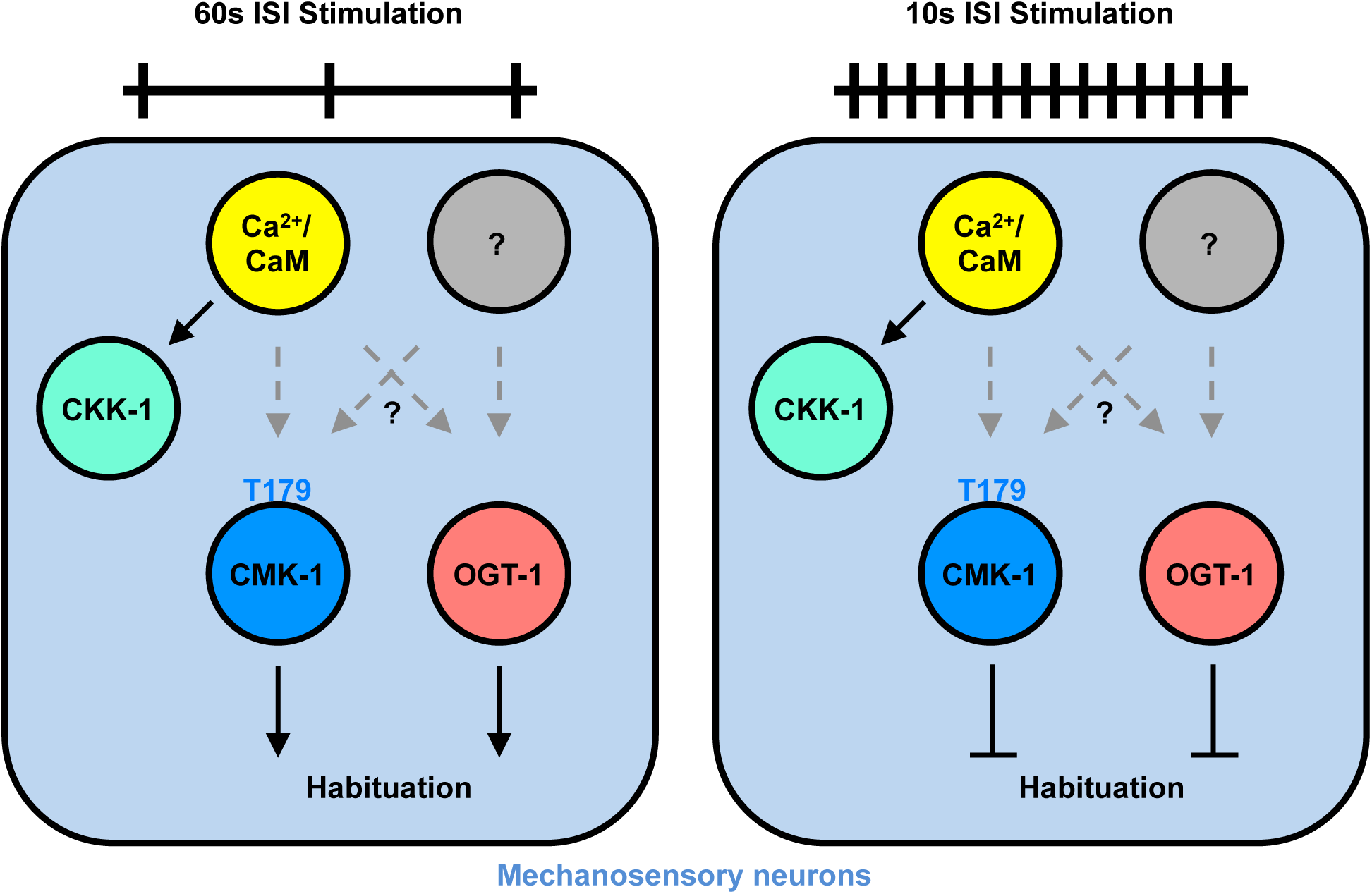
CMK-1 and OGT-1 function in parallel pathways to modulate habituation in an ISI-dependent manner.

It is important to note that the 60s ISI tap habituation phenotype of day 2 adult *cmk-1* mutants could be interpreted as hyper-responsivity to stimuli throughout the training session, rather than an alteration in habituation – which is the ability of animals to learn to decrement their responding to repeated stimulation (Rankin et al. 2009; McDiarmid et al., 2017). However, several lines of evidence from this work suggest that CMK-1 alters habituation as well as sensitivity. First, when worms were tested at 72 hours post egg-lay (Fig. 1; as opposed to 96 hours), *cmk-1* mutants displayed wild-type initial responses to mechanical stimuli, but a higher final level of responding at a 60s ISI, indicating impaired habituation. Second, 96h old *cmk-1* mutants display a wild-type habituated response size when trained at a 10s ISI (Fig. 2B), whereas a general hyper-responsive phenotype should be present regardless of the stimulus interval used. Third, the *cmk-1(pg58)* mutant displayed larger initial responses than controls, but wild-type asymptotic levels, thereby genetically dissociating these response metrics (Fig. 2F). Taken together, our data suggest that initial and habituated levels of responding to tap stimuli are independent metrics and that CMK-1 modulates habituation in an ISI-dependent manner.

During larval development, the integration of a post-embryonically born mechanosensory neuron (AVM) shifts tap responses from a mix of reversals and accelerations to mostly just reversals. By adulthood, neurons have differentiates and wired into the touch response circuit. However, we previously demonstrated that habituation to taps changed over adulthood, as older animals became less sensitive to changes in stimulus intensity (Timbers et al., 2013). This was not apparent when stimuli were simulated with optogenetic activation of the touch cells, suggesting that the change was occurring at the level of mechanostransduction (Timbers et al., 2013). Unlike wild-type, *cmk-1*(*oy21*) mutants displayed an age-dependent naïve response phenotype, such that their tap reversal size increased as they aged from day 1 to day 2 adults. In wild-type, habituation increased with aging regardless of the ISI. In contrast, the direction of the *cmk-1* habituation deficit was dependent on the ISI. Current work is aimed at understanding the effect of aging on touch cell physiology in *cmk-1* mutants.

While CaMKK has been shown to activate CaMK1/4, CaMK1/4 can also function independently of CaMKK. In *C. elegans,* Kimura *et al*. (Kimura et al. 2002) found that CMK-1 was expressed in more neurons than CKK-1 and although CKK-1 enhanced the CMK-1-dependent phosphorylation of CREB, it was not essential for the process. Similarly, Satterlee *et al*. (Satterlee, Ryu, and Sengupta 2004) found that CMK-1, but not CKK-1, functioned to regulate AFD sensory neuron specific gene expression in *C. elegans.* These findings also appear to be consistent with studies in vertebrate organisms, where CaMK1 is known to have a wider expression profile than CaMKK (reviewed in (Hook and Means 2001)). These data suggest that *(i)* activation of CaMK1/4 by calmodulin may be sufficient to activate the kinase in the context of some biological signaling or *(ii)* CaMK1/4 is activated via another, as yet unidentified protein. Our data suggests that CMK-1 functions independently of its canonical upstream activating kinase CKK-1 to influence behavioural plasticity in this context, yet its phosphorylation site at T179 is still required. This work should stimulate further analysis of alternative CMK-1 activating proteins in other forms of neural and behavioural plasiticity.

We used a kinase substrate prediction algorithm developed by Safaei and colleagues (Safaei et al. 2011) to generate a list of protein candidates predicted to be phosphorylated by CMK-1. Behaviorally screening strains with mutations in our top hits, we found that this list was indeed enriched for genes causing tap response and/or habituation deficits when mutated, as 17/22 strains deviated from wild-type in some aspect of these behaviors. We found that the *C. elegans* O-GlcNAc transferase (OGT) mutants, *ogt-1*, displayed strikingly similar phenotypes to *cmk-1* mutants. O-GlcNAc glycosylation is a unique and dynamic cytosolic and nuclear carbohydrate post-translational modification in which *β*-N-acetylglucosamine is covalently attached to serine or threonine residues of proteins. In contrast to other forms of glycosylation, O-GlcNAc glycosylation occurs rapidly (as quickly as 1-5 min after cellular stimulation; (Song et al. 2008; Golks et al. 2007)) and intracellularly and the sugar is not further modified into complex glycans. Hence, O-GlcNAc glycosylation is thought to be more akin to phosphorylation than to other forms of glycosylation. Both OGT and O-GlcNAcase (the enzyme that removes *β*-N-acetylglucosamine residues; OGA) are highly expressed in the brain (Kreppel, Blomberg, and Hart 1997; Gao et al. 2001) and enriched at synapses within neurons (Cole and Hart 2001; Akimoto et al. 2003).

In *C. elegans,* OGT-1 functions in macronutrient storage (Hanover et al. 2005), dauer formation (Lee et al. 2010), lifespan (Rahman et al. 2010), the glucose stress response (Mondoux et al. 2011), and proteotoxicity in neurodegenerative disease models (Wang et al. 2012). O-GlcNAc glycosylation has been shown to function in long-term memory (Rexach et al. 2012) and in cellular models of plasticity, such as: long-term potentiation (LTP), long-term depression (Din et al. 2010) and paired-pulse facilitation (Tallent et al. 2009). Thus, our evidence of an *in vivo* requirement for OGT-1 in learning in *C. elegans* will very likely be conserved in learning in other species.

Epistasis experiments measuring reversal distance revealed the *cmk-1*; *ogt-1* double mutant phenotype was additive. This suggests that the mutations affect tap responses through disruption of independent pathways. Consistent with this interpretation, detailed analysis *cmk-1* and *ogt-1* loss-of-function phenotypes revealed that the deficits arose through differential effects on reversal speed and duration, which combine to determine overall reversal distance (Figure 7). While the effect of reversal duration was additive for double mutants, *ogt-1* suppressed the increased response speed of *cmk-1* mutants. These data suggest that while *ogt-1* and *cmk-1* function in parallel, they do exhibit some interaction to set response speeds. These data highlight the importance of detailed behavioral quantification in mutant analyses when deciphering genetic interactions underlying complex behavior.

To our knowledge, *cmk-1* and *ogt-1* are among the first *C. elegans* mutants hypersensitive to touch at the behavioral level, despite several pioneering genetic screens for harsh and soft touch mutants (Chen, Cuadros, and Chalfie 2015; Chalfie and Sulston 1981). The *akt-1*;*mfb-1* double mutant has increased mechanoreceptor currents compared to wild-type animals and *mec-19* suppresses the touch response deficit of *mec-4*, but neither increase behavioral responses above wild-type (Chen, Cuadros, and Chalfie 2015; Chen et al. 2016). Furthermore, we have shown that *cmk-1(pg58)* mutants also display mechanosensory hyperesponsivity, suggesting that wild-type shuttling of *cmk-1* between the cytoplasm and the nucleus is necessary for setting response level to mechanical stimuli – similar to its effects on thermotaxis and heat avoidance (Yu et al. 2014; Schild et al. 2014; Kobayashi et al. 2016; Satterlee, Ryu, and Sengupta 2004). Further characterization of *cmk-1* interacting proteins, including those predicted through our catalytic site analysis, may identify additional mechanosensory hypersensitive mutants, an as yet largely uncharacterized family of genes regulating mechanosensation.

Early detailed parametric studies on habituation revealed that the nature of the decrement was dependent on the ISI. This led to the hypothesis that the cellular process mediating habituation was similarly dependent on the rate at which stimuli were presented (Rankin and Broster 1992). CMK-1 and OGT-1 slowing habituation at long ISIs and promoting habituation at short ISIs is the first published evidence to support this hypothesis. In conclusion, we show that CMK-1 functions cell autonomously in primary mechanosensory neurons to mediate mechanoresponding and learning. Unlike most forms of behavioural plasticity studied to date, proper nuclear-cytoplasmic shuttling of *cmk-1* is not required for attenuating responses to repeated mechanosensory stimulation. Further, CMK-1 influences short-term habituation independently of its canonical upstream activating kinase CKK-1. We used genome-wide bioinformatic algorithms to predict CMK-1 phosphosites and provided detailed behavioural characterization of 22 strains carrying mutations in top candidates. Among these, OGT-1 displayed ISI-dependent habituation phenotypes strikingly similar to CMK-1. We showed that OGT-1 also functions in the mechanosensory neurons to modulate habituation, but does so independently of CMK-1. Our list of predicted CMK-1 phosphosites will serve as an important resource to streamline identification of CMK-1 phosphorylation targets, as well as other novel regulators of mechanosensory responding and habituation. Overall, our work identifies two conserved genes that function in sensory neurons to modulate habituation in an ISI-dependent manner, providing insight into the molecular mechanism through which animals calibrate plasticity to different rates of sensory stimulation.

## Methods

### Strains and maintenance

Worms were cultured on Nematode Growth Medium (NGM) seeded with *Escherichia coli* (OP50) as described previously (Brenner 1974). The following strains were obtained from the *Caenorhabditis* Genetics Center (University of Minnesota, USA): N2 Bristol, PY1589 *cmk-1(oy21),* VC691 *ckk-1(ok1033),* RB1468 *dkf-2(ok1704),* VC567 *arf-1.2(ok796)*, VC127 *pkc-2(ok328)*, KG532 *kin-2(ce179)*, RB918 *acr-16(ok789)*, RB818 *hum-1(ok634)*, RB781 *pkc-1(ok563)*, RB1447 *chd-3(ok1651)*, RB830 *epac-1(ok655)*, HA865 *grk-2(rt97)*, NW1700 *plx-2(ev773)*; *him-5(e1490)*, PR678 *tax-4(p678)*, KG744 *pde-4(ce268)*, RB758 *hda-4(ok518)*, RB1625 *par-1(ok2001)*, DA596 *snt-1(ad596)*, XA406 *ncs-1(qa406)*, CB109 *unc-16(e109)*, RB653 *ogt-1(ok430)*, TU3568 *sid-1*(*pk3321*) him-5(*e1490*); *lin-15B*(*n744*); *uIs71*[*Pmyo-2*::mCherry; *Pmec-18*::*sid*-1], BC10002 *dpy-5(e907),* VC40557 (which harbors *cmk-1(gk691866)* among many other mutations (Thompson et al. 2013)) and VG834 *cmk-1(gk691866)*. The following strains were obtained from the National BioResource Project for the nematode (Tokyo Womens Medical Hospital, Japan): FX01046 *ogt-1(tm1046),* FX01282 *T23G5.2(tm1282),* FX03075 *pdhk-2(tm3075),* FX00870 *nhr-6(tm870),* FX04733 *syx-6(tm4733),* FX05136 *R11A8.7(tm5136)*, and FX02653 *rab-30(tm2653)*.

### Transgenic strains

The transgenic *C. elegans* strain VH905 *hdIs30*[P*glr-1*::DsRed2] was a gift from H. Hutter (Simon Fraser University, Canada). The plasmid containing *Pmec-7*::mRFP was a gift from J. Rand (University of Oklahoma Health Sciences Center, USA). The transgenic *C. elegans* strains YT1128 *lin-15(n765); tzEx*[P*ckk-1*::GFP; *lin-15*(+)] and YT2016 *tzIs2*[P*cmk-1*::GFP; *rol-6(su1006)*] and plasmids containing *cmk-1* cDNA were gifts from Y. Kimura (Mitsubishi Kagaku Institute of Life Sciences, Japan) and D. Glauser (University of Fribourg, Switzerland).

The following strains were created for this work: VG183 *yvEx64*[P*cmk-1*::GFP; P*mec-7*::mRFP], VG12 *hdIs30*[P*glr-1*::DsRed2]; *tzIs2*[P*cmk-1*::GFP; *rol-6*(*su1006*)], VG19 *tzEx*[P*ckk-1*::GFP; *lin-15*(+)]; *hdIs30*[P*glr-1*::DsRed2], VG92 *cmk -1(oy21)*; yvEx49[P*cmk-1*::CMK-1; P*myo-2*::GFP], VG100 *cmk -1(oy21)*; *yvEx57*[P*cmk-1*::CMK-1; P*myo-2*::GFP], VG260 *yvEx73*[P*ogt-1*::GFP; P*mec-7*::RFP; *rol-6(su1006)*], VG214 *yvEx70*[P*ogt-1*::GFP; *rol-6(su1006)*] and VG261 *yvEx74*[P*ogt-1*::GFP; P*mec-7*::RFP; *rol-6(su1006)*], VG271 *cmk-1(oy21); dpy-5(e907),* VG279 *cmk-1(gk691866); dpy-5(e907),* VG245 *cmk-1(oy21); ogt-1(ok430),* VG708 *cmk -1(oy21)*; *yvEx*707[P*mec-3*::CMK-1::SL2::GFP; P*unc-122p::*RFP], VG709 *cmk -1(oy21)*; *yvEx*707[P*mec-3*::CMK-1::SL2::GFP; P*unc-122p::*RFP]

### PCR fusion construct primers

The primer sequences for the PCR fusion construct P*cmk-1*::GFP were a gift from D. Baillie (Simon Fraser University, Canada). The forward and reverse primer sequences used to amplify the *cmk-1* promoter were TATCCAAAATCTTGCCGAAAGTA and agtcgacctgcaggcatgcaagctTAAAAAGGGGGATTGGGC, respectively. The forward and reverse primer sequences used to amplify GFP were AGCTTGCATGCCTGCAGGTCGACT and AAGGGCCCGTACGGCCGACTAGTAGG, respectively.

The forward and reverse primer sequences used to amplify the promoter-GFP fusion construct were AGAATGCCGTATCATAAGCGTAA and GGAAACAGTTATGTTTGGTATATTGGG, respectively.

The forward and reverse primer sequences used to amplify the *ogt-1* promoter were CTGTTTTCGATTTGATTCTTCAATCAC and agtcgacctgcaggcatgcaagctCTTCTCGATCGTCTAATCCATTCG, respectively.

The forward and reverse primer sequences used to amplify GFP were the same as those used for P*cmk-1*::GFP (see above).

The forward and reverse primer sequences used to amplify the promoter-GFP fusion construct were CGGTTCGCCTTTTATTATGTG and GGAAACAGTTATGTTTGGTATATTGGG, respectively.

### Imaging procedures

Adult worms were anesthetized on glass microscope slides in 100 mM NaN_3_ dissolved in M9 buffer containing sephadex beads (G-150-50, Sigma-Aldrich, St. Louis, MO) and covered with a 1.5 coverslip. An Olympus Fluoview 1000 Confocal microscope and 60x oil immersion lens was used for imaging. Step size was 0.5 µm. GFP was excited using a 488 nm wavelength laser with emitted light collected through a 491-515 nm bandpass filter. dsRed and mRFP were excited using a 543 nm wavelength laser with light collected through a 600-630 nm bandpass filter. Final figures were generated using Image J (National Institutes of Health, Bethesda, MD) and Adobe Photoshop 7.0 (Adobe Systems, San Jose, CA). *Behavioral testing of mutant strains.* Worms were synchronized for behavioral testing on Petri plates containing Nematode Growth Media (NGM) seeded with 50 µl of OP50 liquid culture 12-24 hours before use. Five gravid adults were picked to plates and allowed to lay eggs for 3-4 hours before removal. The animals were maintained in a 20°C incubator for 96 hours (unless otherwise stated). Plates of worms were placed into the tapping apparatus and after a 100s acclimatization period, 30 taps were administered at either a 60s or a 10s ISI. Any statistical comparisons were carried out on plates assayed concurrently (i.e. on the same day).

For CMK-1 rescue strains, twelve hours prior to testing, 40-60 worms carrying the selection marker were transferred using a platinum pick to a fresh NGM plate. Plates were seeded with 50 µl of OP50 liquid culture 12-24 hours before use.

### Complementation test

Wild-type or *cmk-1(oy21)* males were mated with *dpy-5(e907)* hermaphrodites homozygous for one of the three *cmk-1* alleles (wild-type, *oy21*, or *gk691866*). Tap habituation behavior of non-*dpy* F1 progeny was evaluated.

### Behavioral scoring and statistical analysis

Stimulus delivery and image acquisition was done with the Multi-Worm Tracker (version 1.2.0.2) (Swierczek et al. 2011) and offline data analysis was performed with Choreography analysis software (version 1.3.0_r1035) (Swierczek et al. 2011) as described previously (Timbers et al. 2013). ‘Initial’ comprises reversal responses to the first stimulus, while ‘final’ comprises reversal responses to the final four stimuli. To calculate *τ*(i.e. the number of stimuli required for the response to decay by a factor of 1-1/ e towards asymptote), each plate was fit with a two-term exponential curve, with asymptote estimated as the value of the fit at stimulus 30. Responses from all plates were pooled and metrics were compared across strains with ANOVA and Tukey honestly significant difference (HSD) tests. For all statistical tests an alpha value of 0.05 was used to determine significance.

### Kinase and phosphosite prediction and evolutionary analyses

The kinase substrate specificity prediction matrices (KSSPM) for *C. elegans* CMK-1, human CaMK1 isoforms and human CaMK4 were generated using an updated version of the algorithm originally described in Safaei *et al*. (Safaei et al. 2011). The *C. elegans* CMK-1 KSSPM was used to score all of the hypothetical peptides surrounding each of the serine and threonine residues in the 20,470 known *C. elegans* protein sequences. The top 597 scoring phosphopeptides were examined for their conservation in humans using the algorithm described in Safaei *et al*. (Safaei et al. 2011). The identified human phosphosites were then scored with the KSSPMs for human CaMK1 isoforms and human CaMK4.

### RNAi

NGM agar RNAi plates with 1 mM IPTG were seeded two days before use with overnight liquid culture of *E. coli* strain HT115 carrying the empty vector control or the RNAi vector expressing dsRNA against *mec-4* or *ogt-1* (*ogt-1* clone K04G7.3, ORFeome V1.1 obtained from Open Biosystems). TU3568 adults were bleached onto the RNAi plates and second generation adults were tested.

All data, strains, and reagents available upon request.

## Acknowledgements

We would like to thank Angela Leong, Jing Xu, and Savannah Nijeboer for help running experiments and Andrew C. Giles for useful advice and discussions regarding this research. We would also like to thank Daniel G. Taub and Dr. Christopher Gabel for sharing their OGT-1 findings and RNAi protocols. This work was supported by a Natural Sciences and Engineering Research Council Alexander Graham Bell Canada Graduate Scholarship to ELA and TAT, a Canadian Institutes of Health Research Doctoral Research Award to TAM, and a Natural Sciences and Engineering Research Council Discovery Grant to CHR (NSERC RGPIN 1222216-13).

---

**Figure S1.**
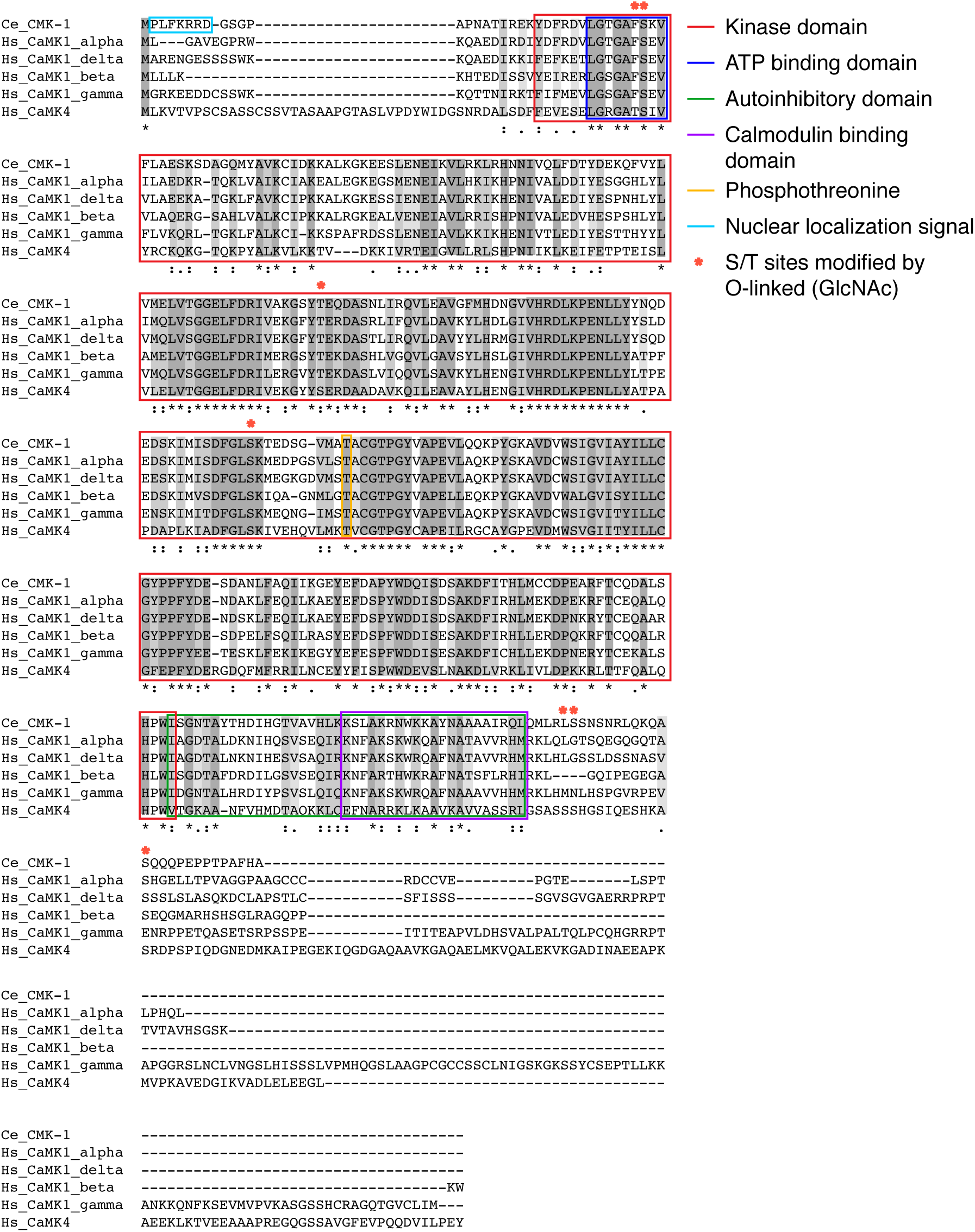
CMK-1 shares homology with human CaMK1’s and CaMK4. Multiple sequence alignment of *C. elegans* CMK-1 (Ce_CMK-1) with human CaMK1*α* (Hs_CaMK1_alpha), CaMK1 *δ* (Hs_CaMK1_delta), CaMK1 *β* (Hs_CaMK1_beta), CaMK1 *γ* (Hs_CaMK1_gamma), and CaMK4 (Hs_CaMK4) highlighting domains and amino acids for phosphorylation and O-GlcNAc modification. Alignment was performed by L-INS-I MAFFT using BLOSUM62 as a scoring matrix and a gap onset penalty of 1.53.

